# Neural responses to social cues in the accessory olfactory bulb are altered by context and experience

**DOI:** 10.1101/2023.11.28.569126

**Authors:** Joseph Dwyer, Maxwell Weinberg, Sarah Y. Dickinson, Joseph Bergan

## Abstract

Social interaction enhances evolutionary fitness by enabling efficient communication of physiological information between individuals. Semiochemicals, that convey socially relevant physiological information, are detected by the vomeronasal organ (VNO) which projects directly to the accessory olfactory bulb (AOB). Mitral and tufted (M/T) neurons in the AOB convey this information from the AOB to a network of brain regions particularly devoted to processing social information and affecting social behavior. The dynamics of social behaviors are shaped by both context and experience. However, our understanding of how alterations in behavior, triggered by the same social cues, correlate with moment-to-moment fluctuations in neural activity within social circuits remains limited. Here, we investigate how context and experience alter the sensory-driven activity of AOB M/T neurons using fiberphotometry and find that the context in which a stimulus is presented can be as important for determining the strength of response as the identity of the stimulus itself.

## Introduction

Social interactions allow animals to increase their evolutionary fitness by communicating physiological information within their own species as well as between species (Green & Marler, 1979; Tinbergen, 1951). Social recognition is necessary for all social behaviors and, in accord with the collective importance of social behaviors, is supported by a dedicated group of brain regions called the social behavior network (SBN; Newman, 1999). The SBN is a deeply interconnected network of brain regions and the activities of neurons distributed throughout this network regulate core aspects of social interactions in all vertebrate species (see Goodson, 2005). In mice, social information depends heavily on a suite of semiochemicals, or pheromones, detected by the VNO and conveyed to central brain regions by the AOB (Chamero et al., 2007; Stowers et al., 2002; Dulac & Torello, 2003; Wysocki et al., 1980; Wysocki & Lepri, 1991; Isogai et al., 2018; Leinders-Zufall et al., 2000; Leypold et al., 2002; Del Punta et al., 2002; Wagner et al., 2006).

Our understanding of pheromonal signaling is most mature for invertebrate species (Kurtovic, 2007; Demir & Dickson, 2005; Herre et al., 2022). However, progress towards identifying the nature and role of mammalian pheromones is accelerating. Recent work performed in rodents has identified import roles for several pheromone candidates: Major urinary proteins produced by the male mouse are both necessary and sufficient to elicit male-male aggression (Chamero et al., 2007; Kaur et al., 2014; Schoeller et al., 2016); Mating is elicited and regulated by methanethiol, Darcin, and ESP1 (Lin et al., 2005; Roberts et al., 2010; Ishii et al., 2017); And formyl peptide receptor-like proteins detect pathogens and identify sickness in other mice (Rivière et al., 2009). Collectively, these results support a labelled line organization in which sensory cues, linked by designated neural circuits, drive specific social behaviors.

While social behaviors are clearly driven by specific sensory cues, they also depend on the context in which a social cue is received (Kaur et al., 2014; Oram and Card, 2022; Zhao, et al., 2021). For example, shoaling fish from high predation environments consistently form larger, more cohesive, shoals compared to similar groups from lower predation environments (Herbert-Read et al., 2017). The presence of a predator, similarly, promotes grouping behaviors and decreases feeding in mammals, while hunger suppresses social interactions in support of food seeking (Creel et al., 2014; St-Cyr et al., 2018; Padilla et al., 2016). Mating and aggressive behaviors are also impacted by the novelty or familiarity of a given social partner (Beach & Jordan, 1956; Bruce, 1959; Davitz & Mason, 1955; Dewsbury, D.A., 1981; Wilson, et al., 1963; Gao et al., 2017) and past experience with a social partner can persistently alter subsequent sensory driven responses in AOB neurons (Yoles-Frenkel et al., 2022).

We targeted fiber photometry recordings to M/T neurons of the AOB because AOB M/T neurons comprise the projection from the AOB to the SBN where they convey critical aspects of individual identity including sex, age and reproductive status of a social partner as well as the presence of allospecific competitors and predators (Lehman, et al., 1980; Halpern, 1987; Meredith, 1998; Scalia & Winans, 1975; Luo et al., 2003; Ben-Shaul et al., 2010; Bergan et al., 2014; Isogai et al., 2018). Here, we investigate how the context of a social stimulus influences AOB neural responses from two perspectives: 1) How the presence or absence of a predator alters neural responses to social cues in the AOB; 2) How repeated interactions with the same stimulus impact the representation of social stimuli in the AOB. To do this, we developed a novel behavior paradigm consisting of automated, acute, and repeatable presentations of social stimuli and paired this behavior paradigm with *in vivo* fiberphotometry recordings from AOB M/T neurons. Our results show that chemosensory driven responses in AOB mitral and tufted neurons are dramatically altered by both the novelty of a stimulus and context in which a stimulus is presented.

## Results

### Stimulus-evoked AOB mitral and tufted neural responses and corresponding behavior

We recorded neural activity from AOB M/T neurons in 16 mice during social interactions (Figure 1; 9 females; 7 males). Multiple social stimuli were presented in an automated and randomized fashion, and behavior states were scored throughout the experiment as exploratory (no engagement with a social stimulus) or social interaction (engaged with an available social stimulus; Figure 2). Experimental mice moved freely throughout the arena and the percentage of time a mouse spent near each stimulus increased sharply when that stimulus was available for social interaction (Figure 2C, D). We found that experimental animals investigated each presented stimulus more than 80% of the time the stimulus was available (Figure 2E). Stimuli were presented in two different configurations (male, female, control) and (male, female, predator). The rate of successful interactions was higher, all around, for stimulus sets that included predator, but in both cases was maintained at a constant level for the duration of the experiment (Figure 2F).

**Figure 1.**
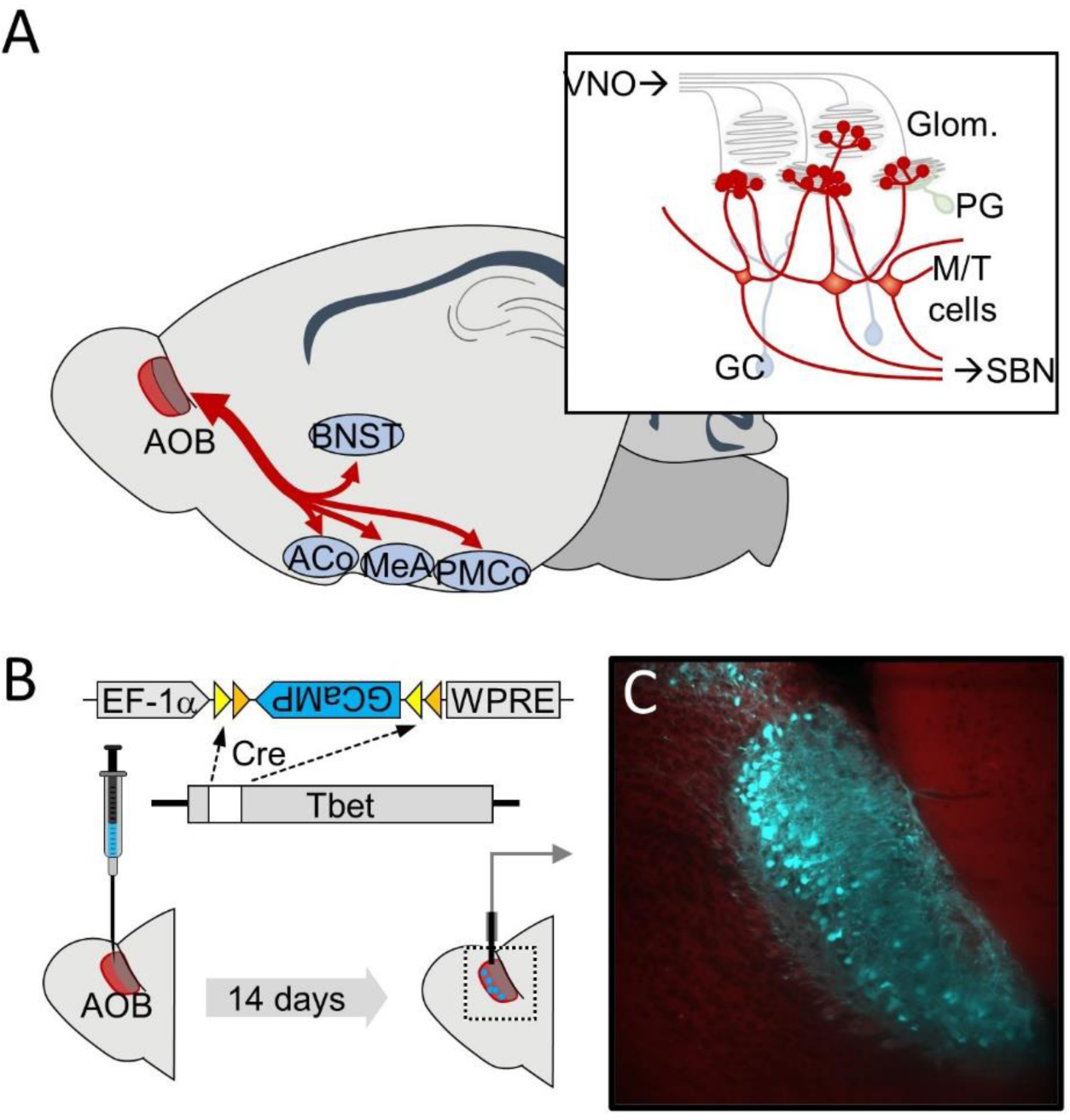
Transgenic and viral strategy to record from AOB projection neurons during social interactions. (A) The AOB receives input directly from the VNO and makes, largely reciprocal, connections with subcortical brain regions devoted to social behavior (red arrows; BNST: bed nucleus of the stria terminalis; ACo: cortical amygdaloid nucleus; MeA: medial amygdala; PMCo: posteromedial cortical amygdala). (Inset) Mitral and tufted neurons are the projection neurons from AOB to the social behavior network (SBN; Glom: glomerular layer; PG: periglomerular cells; M/T: mitral and tufted cells; GC: granule cells). (B) Fiberphometry recordings from AOB mitral and tufted cells rely on conditional AAV-driven expression of GCaMP in a Tbet:Cre mouse paired with a fiberoptic probe implanted in the AOB (Haddad, et al., 2013). (C) Histology showing accurate GCaMP expression (cyan) in M/T cells.

**Figure 2.**
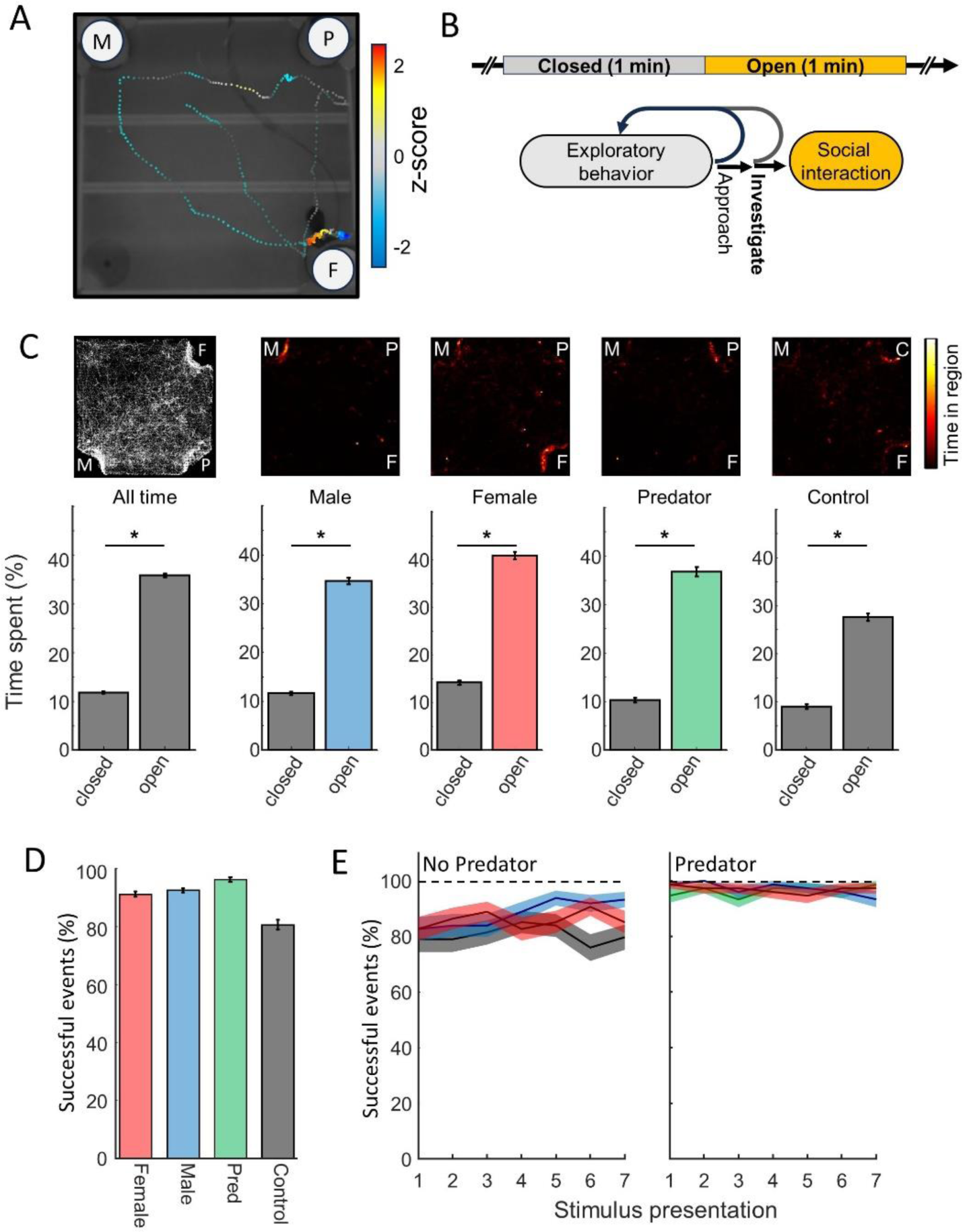
Strategy for automating social interactions during fiberphotometric recording. (A) Trace of mouse position of over 30 seconds (color: intensity of AOB GCaMP signal; M: male; F: female; P: predator; C: control). (B) Social stimuli were presented once every 2 minutes for a 1-minute duration and behavior states were manually scored (exploratory behavior, social interaction, approach, and investigation). (C) Top: density plots indicating the location of an animal during the time a stimulus is available (Left; all time; center left: male stimulus; center: female; center right: predator; right; empty cup). Bottom: the average time (percent of total) spent investigating a stimulus when the stimulus is available (open) or not available (closed; asterisks indicate p<0.0001, paired t-test). (D) Percentage of stimulus presentations that resulted in a successful investigation by the experimental animal (error bars: SEM). (E) Fraction of stimulus presentation events that resulted in a successful investigation event for 7 successive presentations of each stimulus (3 different randomly interleaved stimuli; 21 total) when there was not a predator (left) or was a predator (right) in the stimulus set (red: female; blue: male; green: predator; grey: empty cup; shaded regions: SEM).

Animals explored the arena broadly when a stimulus was not available and then typically approached the stimulus rapidly when it became available (Figures 2A, 3A). AOB M/T activity sharply increased after the experimental animal initiated an investigation of the stimulus animal and the increase in GCaMP activity was sustained above baseline for the duration of the interaction (Figure 3B, C). The rapid increase in GCaMP activity was often preceded by a transient decrease in AOB M/T activity that began immediately before the start of the investigation as can be seen in the response to male stimuli (Figure 3C). Across the population, strong sensory responses were observed in AOB M/T neurons to male stimuli (Mean: 0.19 +/- 0.02 SEM; p-value<0.000001) and female social stimuli (Figure 4; Mean: 0.21 +/- 0.03 SEM; p-value<0.000001) with no significant sex differences observed in how AOB neurons in male versus female mice responded to social stimuli (female stimuli: p-value=0.58; male stimuli: p-value=0.13). Clear responses were also observed in response to predator stimuli (Mean: 0.18 +/- 0.03 SEM; p-value<0.000001), while the average response to control stimuli was not significantly different from the baseline level of activity in AOB M/T neurons (Mean: 0.03 +/- 0.03 SEM; p-value: 0.28963)

**Figure 3.**
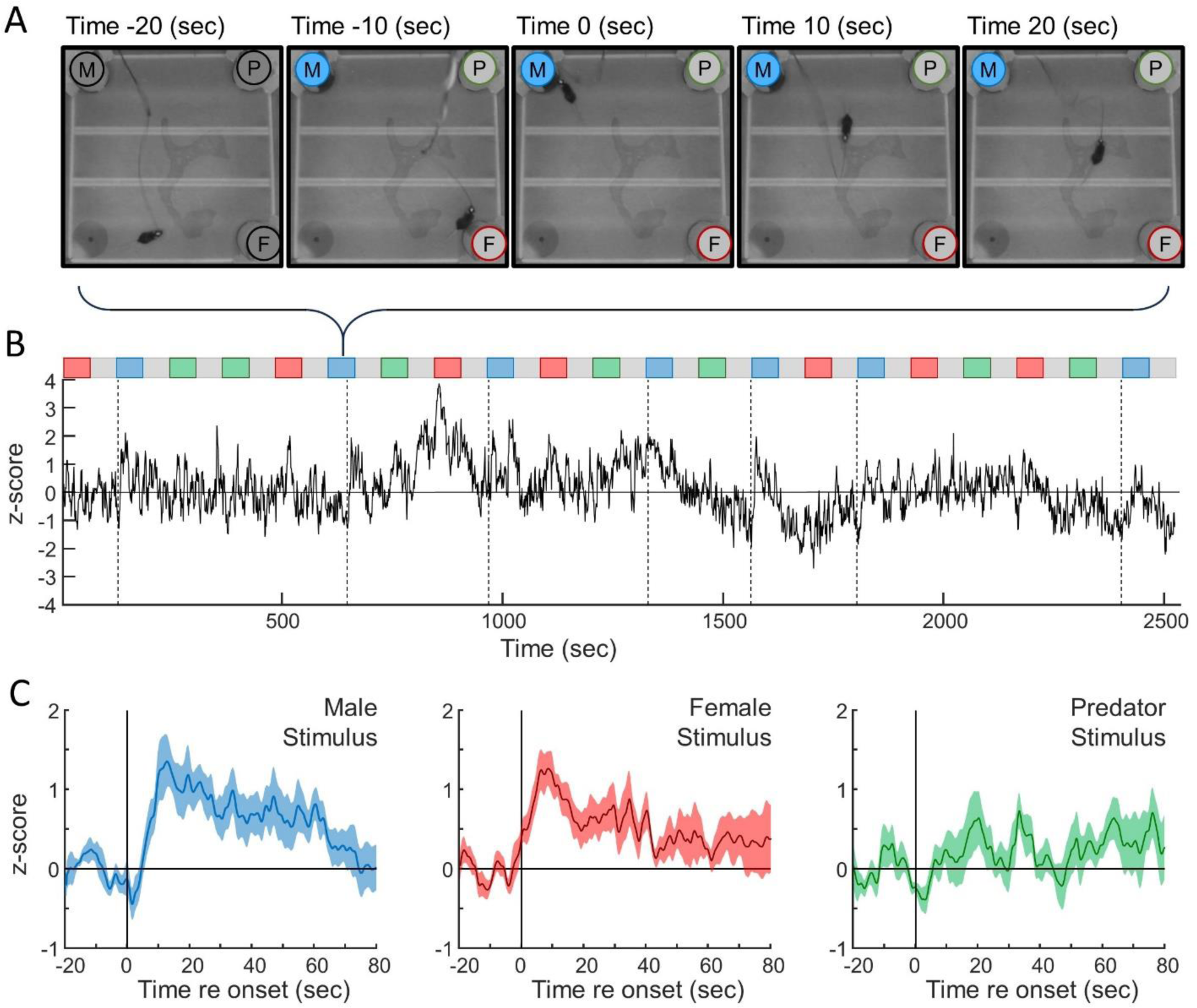
Example of sensory responses of AOB neurons to social and predator stimuli in an adult female mouse. (A) Video frames surrounding a social interaction, with 0 seconds (center) aligned to the onset of stimulus investigation (Blue shading indicates the availability of the male stimulus). The predator (P) and female (F) stimuli were closed during the duration of the shown frames (bracket indicates the investigation event in panel B highlighted in panel A). (B) GCaMP signal (black line; z-score normalized) during seven randomized presentations of three stimuli (male, female, predator; order indicated by the colored boxes). Vertical dashed lines indicate the start of all investigations of the male stimulus. (C) Average responses (from panel B) aligned to the start of investigation of male, female, and predator stimuli (shading: SEM).

**Figure 4.**
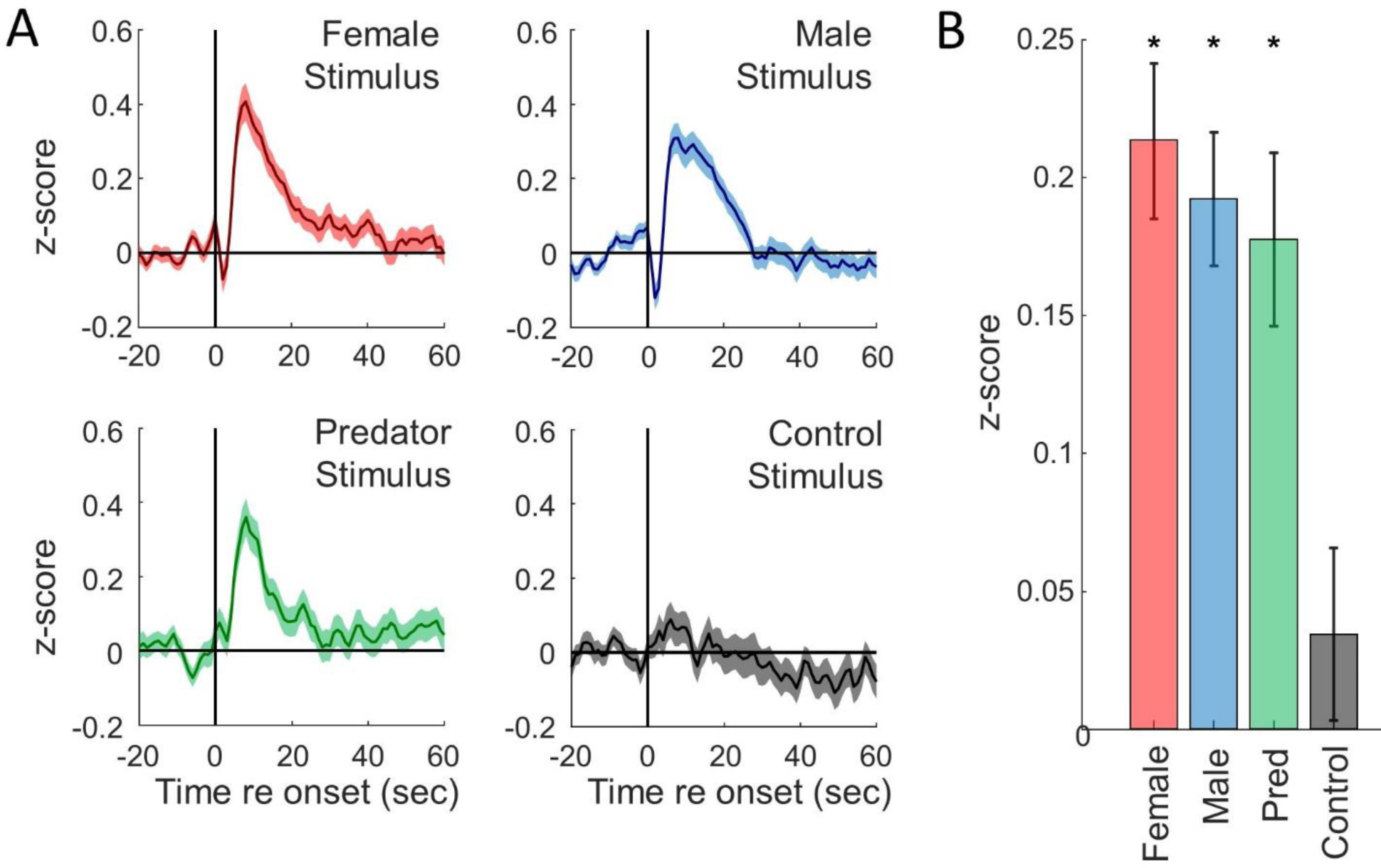
Average GCaMP response across stimulus presentations. (A) Average GCaMP signal of female (red), male (blue), predator (green) and control (grey) stimuli relative to the first investigation after a stimulus became available (shading: SEM; N = 16: 7 males, 9 females). (B). Average z-score during the 20 seconds following sensory investigation for female, male, predator, and control stimuli. Error bars: SEM; asterisks: p-value<0.000001.

### AOB M/T neurons respond differentially to social stimuli in the presence of predator stimuli

Inclusion of predator stimulus increased AOB responses to social stimuli compared to responses during experiments that omitted predator stimuli (Figure 5). Importantly, predator stimuli were never presented simultaneously with social stimuli; but rather, influenced responses to subsequent stimulus presentations and social behavior. Inclusion of a predator stimulus roughly doubled the AOB response to female (Figure 5A; female response with predator: 0.29 +- 0.04; female response without predator: 0.15 +- 0.04; p-value=0.01) and male stimuli (Figure 5D; male response with predator: 0.25 +- 0.04; male response without predator: 0.14 +- 0.03; p-value=0.018)). The sensory-driven AOB response was strongest to the first stimulus presentation and declined with subsequent presentations of the same stimulus (Figure 5B, C) and was enhanced in the presence of a predator for all stimulus presentation trials (Figure 5C,F). In contrast to the decrease in sensory-driven response in successive stimulus interactions, the level of investigatory behaviors stayed steady throughout the duration of the experiment for male, but not female experimental animals. Female experimental animals displayed greater investigation of male stimuli when a predator was included in the stimulus set, while no difference was observed for females investigating females nor did the presence of a predator in the stimulus set impact the time males spent investigating either male or female stimuli (Figure 2 Supplement 1).

**Figure 5.**
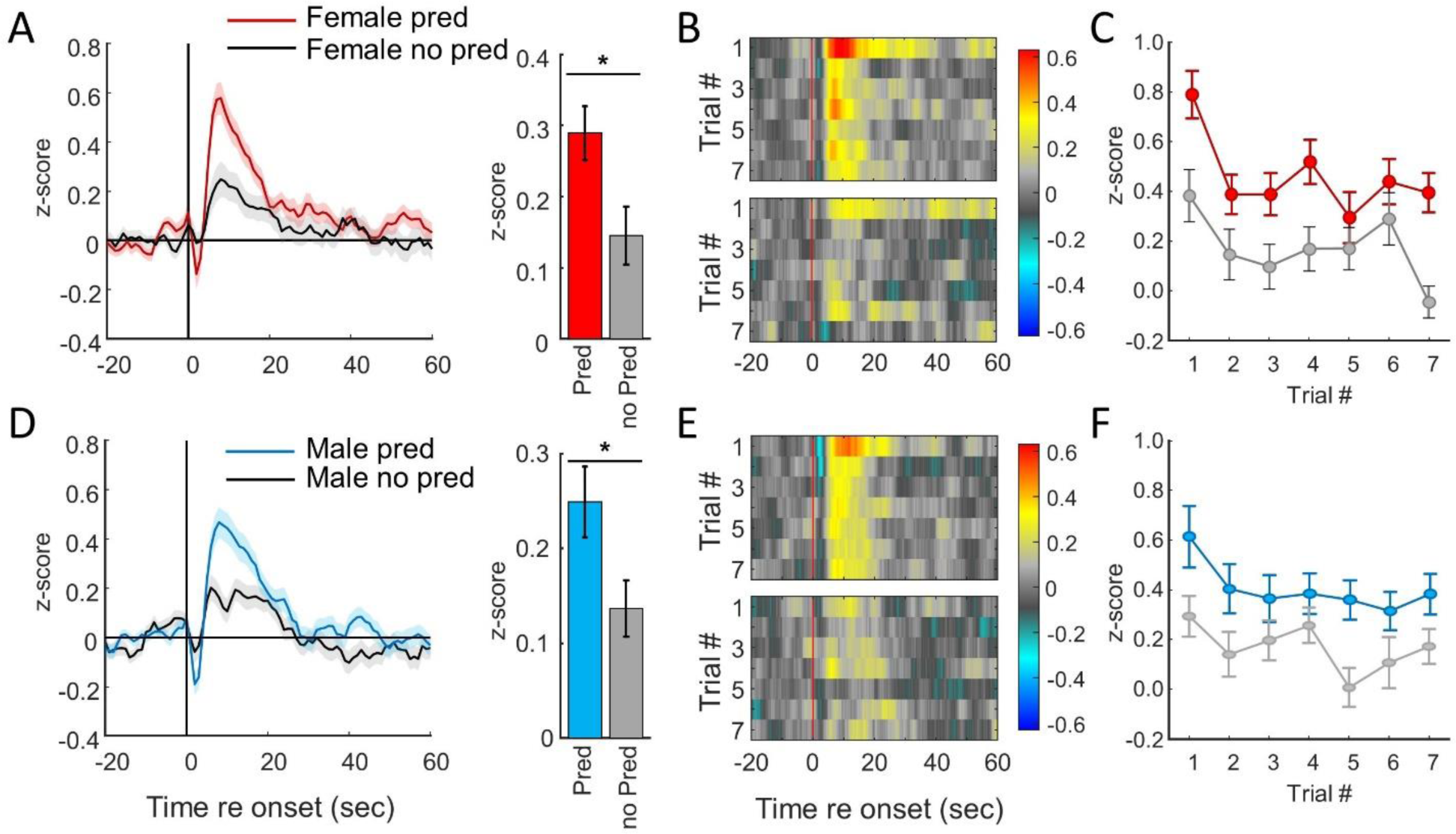
AOB response to social stimuli depends on context. (A) Left, average GCaMP signal (z-score normalized) across presentations of female stimuli during experiments that included predator stimuli (red) or did not include predator stimuli (grey). Right, mean GCaMP response to female stimuli during predator and no predator experiments (t-test comparison p value = 0.011, error bars: SEM). (B) Pseudocolored plot of the average response to female stimuli for each of 7 presentations during the predator condition (top) or no predator condition (bottom). (C) Effect of repeated presentations of female stimuli on the average GCaMP response (0-20 seconds) for predator (red) and no predator (gray; error bars: SEM). (D) Left, average GCaMP signal (z-score normalized) across presentations of male stimuli during experiments that included predator stimuli (blue) or did not include predator stimuli (grey). Right, mean GCaMP response to male stimuli during predator and no predator experiments (t-test comparison p value = 0.018; error bars: SEM). (E) Pseudocolored plot of the average response to male stimuli for each of 7 presentations during the predator condition (top) or no predator condition (bottom). (F) Effect of repeated presentations of male stimuli on the average GCaMP response (0-20 seconds) for predator (blue) and no predator (gray; error bars: SEM).

### AOB neural responses to novelty and habituation

We presented social stimuli either rarely or commonly to determine how familiarity with a stimulus animal impacted sensory responses of AOB M/T neurons. Stimulus A was presented six consecutive times followed by a single presentation of stimulus B (age and sex matched to stimulus A). A fixed order of seven stimulus presentations (6: A; 1: B) was repeated 3 times for a total of 21 stimulus presentations during each experiment (Figure 6A). M/T responses decreased with successive presentations of either stimulus A or stimulus B (Figure 6B-E). Responses to stimulus A decreased dramatically with contiguous stimulus presentations while the response to stimulus B, with breaks between each successive presentation, were maintained at a consistently high level (Figure 6C, D). The less frequently presented stimulus B resulted in stronger AOB neural activity, when presented, than the more frequently presented stimulus A (Figure 6C; paired t-test, p = 0.02). The first presentation of stimulus A (trial 1) produced the strongest overall response, and the first presentation of stimulus B (trial 7) produced the second strongest overall response (Figure 6B, D). Across the population, responses to novel stimuli (first presentation of a stimulus) were roughly twice as strong as observed for familiar stimuli (Figure 6E; paired t-test, p < 0.00001).

**Figure 6.**
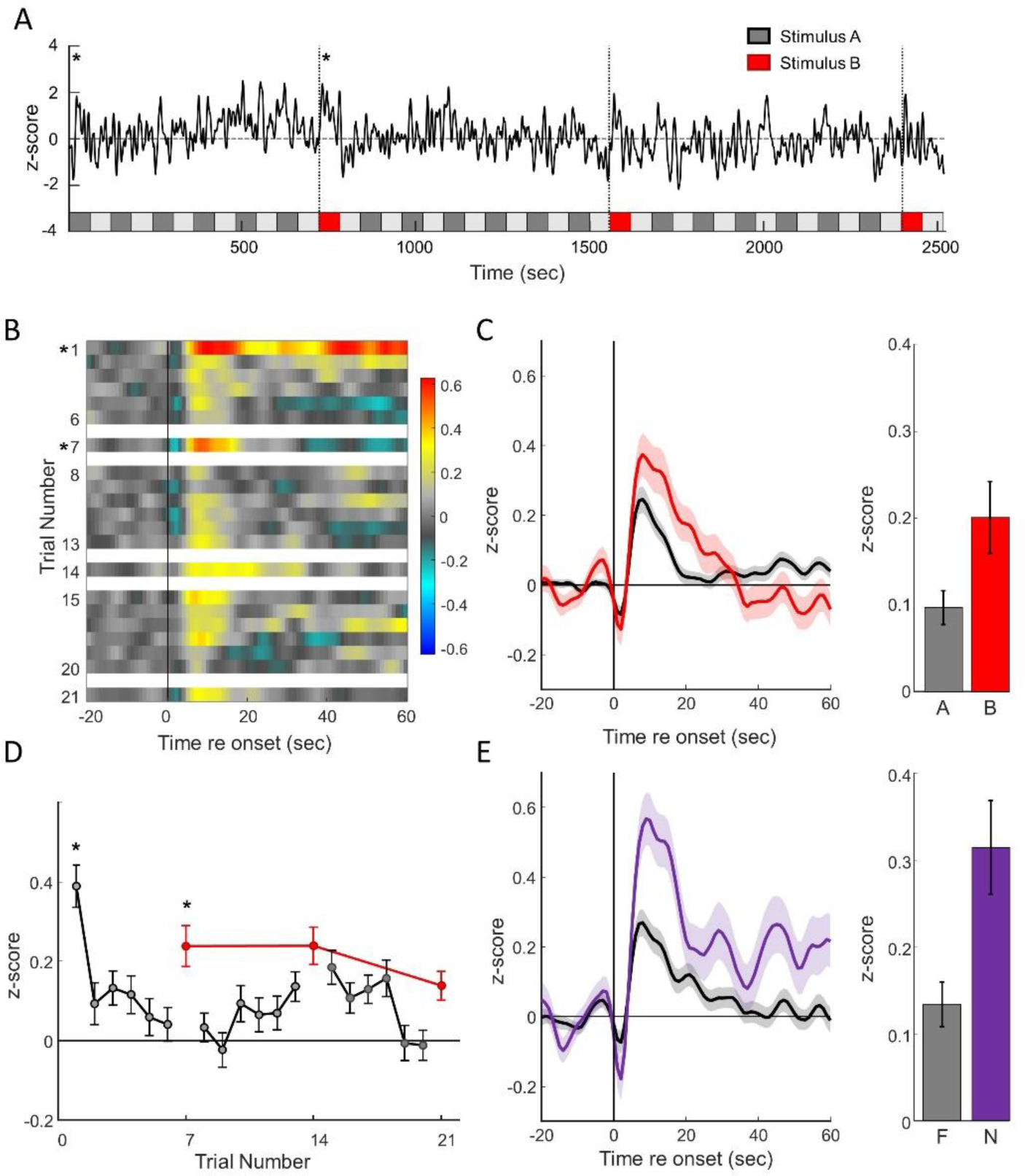
AOB responses convey stimulus novelty on multiple time scales. (A) GCaMP signal (black line; z-score normalized) during an initial baseline period followed by 18 presentations of stimulus A and 3 presentations of stimulus B presented in a standardized order shown by the boxes below. Vertical dashed red lines indicate investigation events to the less frequently presented stimulus. (B) Pseudocolor plot representing the average GCaMP signal aligned to the start of a sensory investigation. The stimulus order is as shown in panel A and changes between the presented stimulus are represented by white space. Purple asterisks represent novel trials. (C) Left, average GCaMP signal across presentations to rare (stimulus B: red) or frequent (stimulus A: gray) age and sex matched social stimuli. Right, mean GCaMP response to rare stimuli and frequent stimuli. (D) Average AOB GCaMP response to frequent (grey) and rare (red) stimuli during the 20 seconds following stimulus investigation (error bars: SEM). (E) Left, average GCaMP signal across presentations to novel (first presentation of either stimulus A or B: purple; asterisks in A,B, D) or familiar (subsequent presentations of either stimulus A or B: gray) age and sex matched social stimuli. Right, mean GCaMP response to novel (N) stimuli and familiar (F) stimuli.

## Discussion

Natural social behaviors typically progress from an initial investigation towards consummatory behaviors such as aggression, parenting, or mating within the span of several minutes. The progression of behaviors during these interactions is critical to understand, and correlating specific behaviors with changes to neural activity is difficult when each social interaction can follow a unique trajectory. In the experiments reported here, social interactions were largely restricted to investigatory behaviors during defined epochs of time and were highly reproducible and consistent. Because social interactions were restricted, the fiberphotometry results presented here are most relevant for understanding the sensory coding during the early stages of social investigation.

The strategy to study social interactions introduced here allowed us to reliably assess neural responses in awake mice during automated and randomized presentations of multiple social stimuli. This approach relies on the innate drive of mice to investigate social and allospecific stimuli. Even without training or previous experience, mice successfully investigated each stimulus more than 80% of the time within the first 60 seconds the stimulus was available. Because each stimulus presentation was temporally precise, it was possible to investigate responses to repeated and consistent social interactions in a short period of time. This approach allowed us to systematically vary an animal’s interactions to quantitatively investigate the effects of both context and previous experience on the activity of AOB M/T neurons. When used in combination with approaches such as a resident intruder assay, this novel approach should help disambiguate neural activity linked to sensory investigation versus neural activity linked to the execution of social behaviors.

The reported fiberphotometry recordings were restricted to the output layer of the AOB (M/T cells) and reflect the chemosensory information conveyed from the olfactory bulb to central brain regions involved in social behaviors (Goodson, 2005; Newmann, et al., 1999; O’Connell and Hofmann, 2012). Consistent with previous studies, AOB M/T neural activity was driven by both social and predator stimuli (Ben-Shaul et al., 2010; Bergan et al., 2014; Ishii et al., 2017; Lin et al., 2005; Luo et al., 2003; Martel & Baum, 2007; Stowers et al., 2002). However, AOB sensory responses depended almost as much on the context in which a stimulus was presented as the identity of the stimulus itself.

The presence of a predator is a salient feature of the environment with profound implications for survival. We initially predicted that a predator stimulus would inhibit subsequent sensory responses to conspecific stimuli. Instead, we found that responses to conspecific stimuli were enhanced when a predator stimulus was included in the stimulus set. Wild mice gather in larger groups, likely to buffer risk factors, when a predator is nearby (Apfelbach, et al., 2005). When a predator stimulus was present in our experiments, both male and female animals more consistently investigated each stimulus (Figure 2C). However, the addition of a predator stimulus did not change the average amount of time male mice spent near either conspecific stimulus, nor how female mice investigated female mice. In contrast, females spent more time investigating a male when a predator stimulus was included. Therefore, the addition of a predator to the stimulus set increased the magnitude of sensory response to conspecific stimuli consistently, while the impact of including a predator stimulus on investigative behavior displayed a clear sex difference. Given that stimuli were presented one at a time, these experiments were not able to distinguish whether the addition of a predator stimulus biases investigative biases towards one social stimulus.

We suggest that the observed difference in AOB responses reflects a different internal state that occurs in the presence of a predator that alters the responsivity of M/T neurons throughout the duration of the experiments. Indeed, the enhanced responses to conspecific stimuli occur at a time when the predator stimulus is not available. Our data do not distinguish potential mechanisms that drive enhanced responsivity when a predator stimulus is included. One possibility is that local interactions, intrinsic to the AOB, shape responses in a context-dependent manner (Castro, et al., 2007). A second possibility is that the enhancement of social responses observed in the presence of predator stimuli is mediated by feedback from central regions involved in social behavior or in regulating intrinsic state (Gupta et al., 2015; Inbar, et al., 2021). A third possibility is that changes in hormone signaling (e.g., cortisol) resulting from predator exposure may influence the strength of AOB responses to social stimuli. These possibilities are not mutually exclusive, and future research is needed to clarify these possibilities.

AOB sensory responses to social stimuli adapted profoundly and rapidly with successive stimulus presentations. The first (novel) stimulus presentation elicited the strongest response and responses fell nearly to baseline levels after only six repetitions. Novel animals can elicit remarkably different behavioral responses than familiar animals and these differences are mediated by the SBN (Wallace, et al., 2023). For example, a novel male mouse produces pregnancy block in a mated female (Bruce, 1959) and a novel conspecific elicits distinctly more investigatory behaviors and social interaction (Dewsbury, D.A., 1981). The rapid habituation of AOB responses to social stimuli is consistent with, and may partially underlie, the changes in social behaviors elicited by familiar stimuli. These experiments establish adaptation and habituation, common features of sensory systems (Ulanovsky & Nelken, 2003; Bergan & Knudsen, 2009), as important considerations for understanding sensory coding in the AOB.

AOB M/T neuron responses to multiple, progressive investigations to the same stimuli habituated dramatically, and presentation of novel stimuli resulted in a stronger response that also increased the subsequent response to the initial social stimulus. This result highlights the importance of novelty and familiarity in guiding social behaviors, and demonstrates that, even as early in the sensory processing pathway as the AOB, neurons have access to how novel or familiar a stimulus is. This novelty-dependent difference in AOB activity may mediate naturally occurring differences in consummatory behaviors directed towards novel versus familiar animals.

These experiments provide new insight into how sensory information is processed by the AOB to, ultimately, generate social behaviors. We isolated the period of behavior most relevant for sensory investigation and presented social stimuli in a reproducible and acute manner. This quantitative approach to investigating the neural basis of social interactions in freely behaving and untrained mice allowed us to identify the large impact of context, including the presence of a predator or familiarity with a social partner, impact sensory responses in the AOB.

## Methods

### Animals

All experiments were performed in strict compliance with the National Institute of Health. All animals were handled according to a protocol approved by the UMass Amherst Institutional Animal Care and Use Committee (IACUC; protocol #2018-0014 and #2017-0060).

Sixteen adult Tbet-cre mice 2 to 8 months old (female: n=9; male: n=7) were housed in a temperature (22°C) and light (12hr: 12hr light: dark) controlled facility, with ad libitum access to food and water. Tbet-cre mice expression of cre faithfully recapitulates endogenous Tbet expression (Haddad, et al., 2013) and display no known behavioral deficits in either heterozygous or homozygous animals.

### Surgeries

We injected a conditional GCaMP expressing AAV virus (AAV9:FLEX:GCaMP6s; Addgene Plasmid #:100845; Chen, et al., 2013) in the AOB (Bregma 3.5, Lateral 1.0, Depth 1.0) of Tbet-cre mice to target M/T cells in the AOB. Fiberoptic implants were custom made by inserting a 10 mm length of 440 µm diameter silica optical fiber (Polymicro, FIP400440480) into a 440 µm ceramic ferrule (Thor Labs, ADAF2) and securing with optical glue. Fibers were implanted into the same location used for initial AAV injections and secured to the skull using dental cement. Animals were left to rest for 72 hours after surgery and were subsequently single housed for the duration of the experiment.

### Fiber Photometry Signal Collection and Processing

Experimental animals were attached via a fiberoptic patch cord to a fiberphotometry system (Doric, RFPS_S). A pig-tailed optical rotary joint allowed animals to freely roam in the box for the extent of all experiments (∼60 minutes). Fiber photometry recordings were made using lock-in sampling with excitation wavelengths of 470 nm for the calcium dependent signal (GCaMP) and 405 nm for the calcium independent signal for control (isosbestic). GCaMP and isosbestic excitation were sinusoidally modulated at frequencies of 572.205 Hz and 208.616 Hz respectively and the GCaMP and isosbestic signal was calculated for these frequencies. The calcium dependent signal was isolated using regression normalization (GCaMP vs isosbestic) and subtracting the scaled calcium-independent (isosbestic) signal from the calcium-dependent GCaMP signal (Figure 2 Supplement 2). All photometry signals were then z-score normalized for further analysis.

### Behavioral Paradigm

Fiberphotometry was synched with behavioral data via an LED, in view of the video recording, that turned on during photometry data collection. Fiber photometry (experimental) animals were placed in a customized stimulus presentation arena (24” by 24”) that contained rotating computer-controlled cages in each corner. Age-matched male or female animals, a clean nestlet (control), or a nestlet soaked with 1 mL of 1:1000 diluted fox urine (predator stimuli) was placed in the cages and presented in a randomized order under computer control. A computerized schedule consisted of a one-minute pre-experiment baseline, followed by seven presentations each of a male, female and control or predator stimulus in pseudorandom order. Each presentation of stimuli consisted of a one-minute presentation period, followed by a one-minute period where no stimulus is presented. For novelty and habituation experiments, two age and sex matched stimulus animals were presented with the same interstimulus timing as control and predator experiments. Stimuli A (familiar) was presented in six repetitions followed by one presentation of stimuli B. This pattern was repeated three times during the experiment for a total of 21 stimulus presentations.

Social and non-social behaviors were manually scored for the duration of the experiment including social interaction, exploratory behavior, investigation, and grooming. Behavior scoring relied on the physical location of the experimental animal within the behavior box and the state (available or unavailable) of the stimuli. Engagement was defined by the experimental animal occupying a small radius around the open cage window with the head directed at the stimulus animal. Stimulus presentations were only included in the analysis if the experimental animal investigated while the stimulus was available and fiberphotometry signals were aligned to the start of the first investigation event. All manual behavior scoring relied on a custom MATLAB script-based interface that allowed behavior classification time locked to fiberphotometry. (Mathworks). We used pose estimation (DeepLabCut, Mathis et al. 2018) to track the position of six points (nose, left and right ears, torso, base of tail, and fiber implant) of the experimental animal during each experiment (Figure 2A).

### Statistical Analyses

We determined the amount of time that an animal spent within 12 cm of an available stimulus, as well as an unavailable stimulus, for all tested stimuli. A repeated samples t-test was performed comparing the amount of time investigating each stimulus and determining the impact of the stimulus, the experimental animal’s sex, the novelty of the stimulus, and the presence or absence of a predator stimulus. Multiple comparisons were Bonferroni corrected.

Statistical analysis of fiberphotometry signals were conducted on a 20 second time window starting with the onset of voluntary stimulus investigation. Fiberphotometry signals were expressed as a z-score and the mean response was calculated for each experiment. A repeated samples t-test was then conducted comparing the mean responses for each experiment using the ‘ttest2’ function in MATLAB (Mathworks).

### Tissue Processing

Following fiberphotometry data collection, animals were deeply anesthetized with isoflurane and perfused with 50ml cold PBS followed by 25 ml cold PFA (4% in PBS). The brain was extracted and post-fixed in 25 ml PFA (4% in PBS) at 4 °C overnight, and then sliced at 100 µm on a vibratome for posthumous verification of implant targeting and GCaMP expression.

**Figure 2 Supplement 1:**
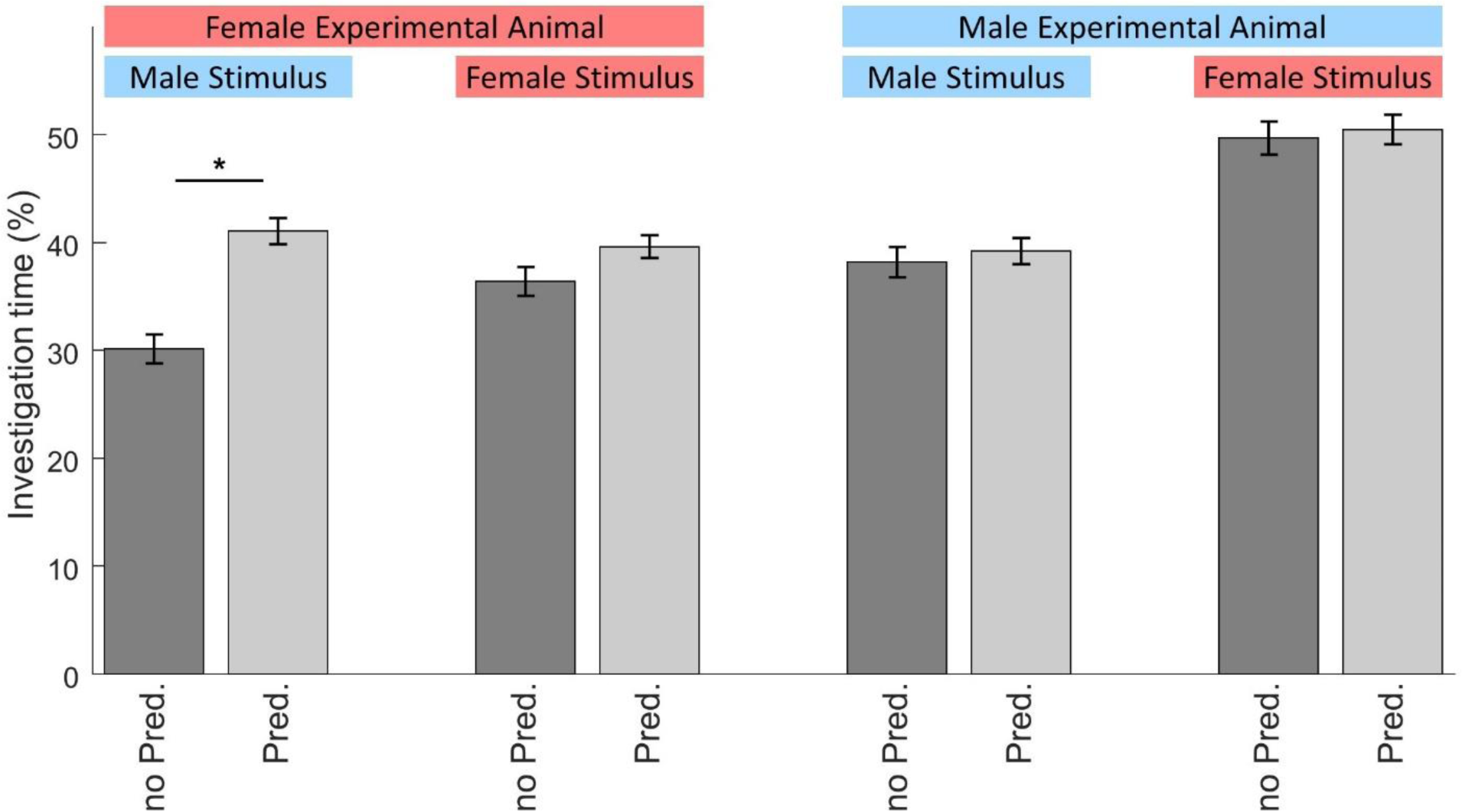
The impact of a predator on social investigation in male and female mice. Investigation time was defined by proximity (12 cm: see methods) to an available stimulus. The percentage of time, of the full time the stimulus was available, is plotted. Error bars: SEM; asterisks: p-value<0.0001 (all other comparisons: p-value> 0.2)

**Figure 2 Supplement 2.**
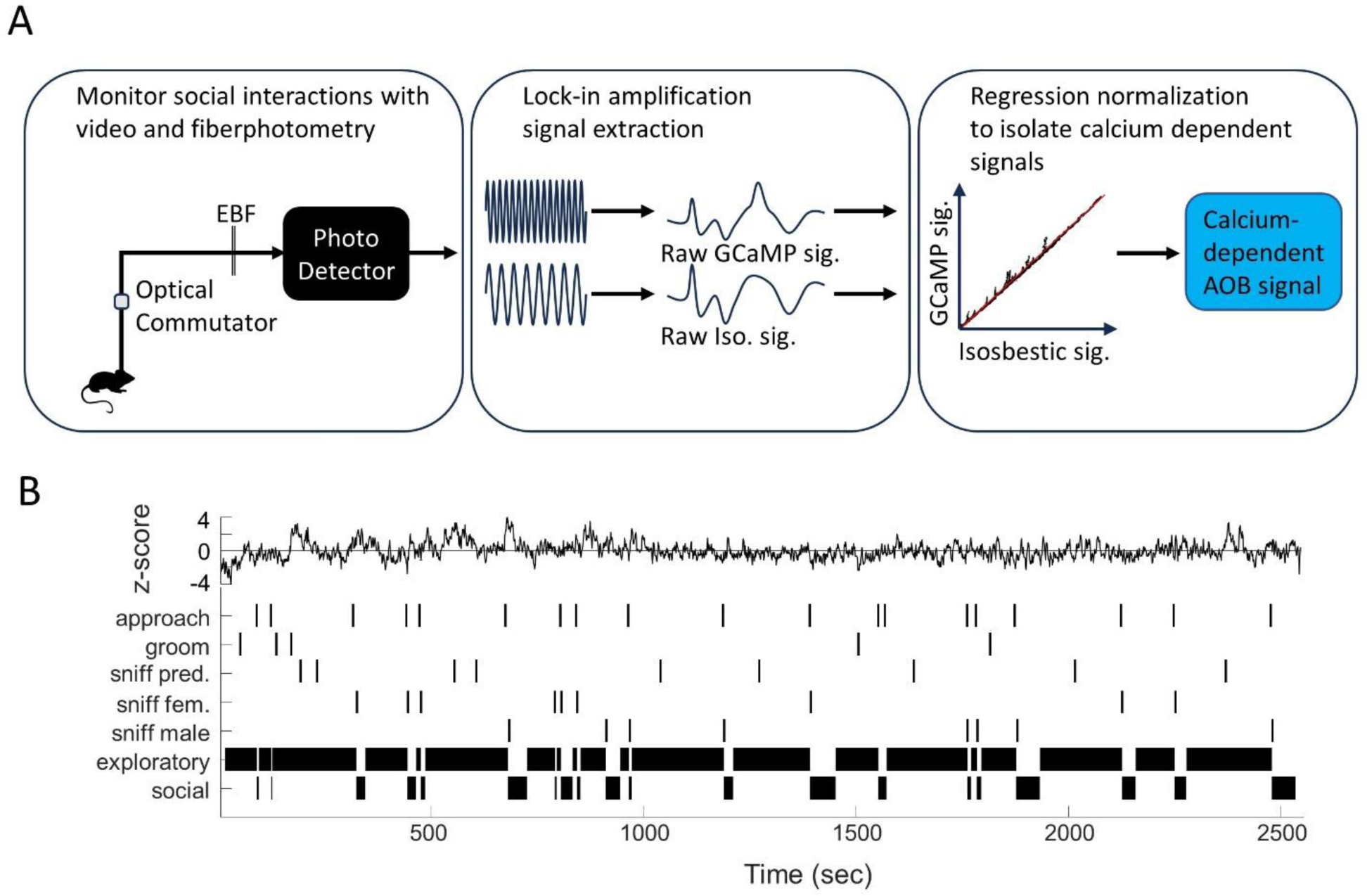
Analysis pipeline for fiber photometry signals and behavior. (A) GCaMP and isosbestic signal are collected and synced with behavior (left). GCaMP and isosbestic signals are separated utilizing lock-in sampling and amplification (middle). GCaMP and isosbestic signals are compared via regression normalization allowing the isolation of calcium dependent AOB signals from movement. (B) Scored behavioral states and events are synced with calcium-dependent AOB signal z-scores.

## Notes

### Competing Interest Statement

The authors have declared no competing interest.

